# Structural basis for +1 ribosomal frameshifting during EF-G-catalyzed translocation

**DOI:** 10.1101/2020.12.29.424751

**Authors:** Gabriel Demo, Anna B. Loveland, Egor Svidritskiy, Howard B. Gamper, Ya-Ming Hou, Andrei A. Korostelev

**Affiliations:** RNA Therapeutics Institute, Department of Biochemistry and Molecular Pharmacology, UMass Medical School, Worcester, MA, USA; Central European Institute of Technology, Masaryk University, Kamenice 5, Brno, 625 00, Czech Republic; Department of Biochemistry and Molecular Biology, Thomas Jefferson University, Philadelphia, PA, USA

## Abstract

Frameshifting of mRNA during translation provides a strategy to expand the coding repertoire of cells and viruses. Where and how in the elongation cycle +1-frameshifting occurs remains poorly understood. We captured six ∼3.5-Å-resolution cryo-EM structures of ribosomal elongation complexes formed with the GTPase elongation factor G (EF-G). Three structures with a +1-frameshifting-prone mRNA reveal that frameshifting takes place during translocation of tRNA and mRNA. Prior to EF-G binding, the pre-translocation complex features an in-frame tRNA-mRNA pairing in the A site. In the partially translocated structure with EF-G, the tRNA shifts to the +1-frame codon near the P site, whereas the freed mRNA base bulges between the P and E sites and stacks on the 16S rRNA nucleotide G926. The ribosome remains frameshifted in the nearly post-translocation state. Our findings demonstrate that the ribosome and EF-G cooperate to induce +1 frameshifting during mRNA translocation.

## INTRODUCTION

To accurately synthesize a protein, the ribosome maintains the mRNA reading frame by decoding and translocating one triplet codon at a time^1^. Concurrent ∼25 Å movement of the mRNA and tRNAs is catalyzed by the conserved translational GTPase EF-G in bacteria (EF2 in archaea and eukaryotes)^2,3^. After formation of a peptide bond, the peptidyl-tRNA and deacylated tRNA move from the A and P sites to the P and E sites, respectively. This translocation requires spontaneous and large-scale (∼10°) inter-subunit rotation of the ribosome^4,5^. Despite pronounced rearrangements of subunits and extensive motions of tRNA and mRNA at each elongation cycle, the ribosome maintains the correct reading frame through hundreds of codons^6,7^.

Nevertheless, change of the reading frame, termed frameshifting, is common in viruses, bacteria and eukaryotes, where it enables the expansion of the coding repertoire and regulation of gene expression^8^. During frameshifting, the translating ribosome switches to an alternative reading frame, either in the forward (+; i.e. skipping one or more mRNA nucleotides) or reverse (–; i.e. re-reading one or more mRNA nucleotides) direction. This work focuses on +1 frameshifting (+1FS), which is important for gene expression in various organisms. For example, in bacteria, it regulates expression of the essential release factor 2^9^,^10^. In eukaryotes, +1FS regulates metabolite-dependent enzyme expression^11^ and leads to pathological expression of huntingtin^12^. +1FS can be amplified by dysregulation of ribosome quality control mechanisms^13,14^, and it is being exploited to synthetically expand the coding repertoire of genomes by inserting non-natural amino acids via a tRNA that can perform +1FS^15^. Because +1FS occurs during the dynamic stage of protein elongation, its molecular mechanism has remained challenging to study.

Here we address this challenge by using cryo-EM to visualize +1FS on one of the most +1FS-prone mRNA sequences in the bacterial genome. The mRNA sequences CC[C/U]-[C/U]^16^ induce +1FS due to imbalances in tRNA concentrations^17,18^, lack of tRNA post-transcriptional modifications^19-21^, or nucleotide insertions in the anticodon loop of tRNAs^22-24^. Under normal growth conditions, CC[C/U]-[C/U] sequences in *E. coli* can induce +1FS up to ∼1%^25^, exceeding the average frequency of spontaneous frameshifting on other sequences by two orders of magnitude^26^. *In vitro*, mRNA CC[C/U]-N (N = A, C, G, U) sequences are even more prone to +1FS, achieving 70% efficiency^21^. The CC[C/U]-N sequences code for proline (Pro) and are decoded by two isoacceptors of tRNA^Pro^ with the anticodon UGG or GGG^21^. The isoacceptor tRNA^Pro^(UGG) is essential for cell growth and has the ability to read all four Pro codons^27^. Studies have proposed that +1FS by tRNA^Pro^(UGG) can occur during one of the three stages of the elongation cycle: (1) decoding of a slippery sequence when the tRNA binds to the ribosomal A site^28-30^; (2) EF-G-catalyzed translocation of the tRNA from an in-frame position at the A site to the +1-frame position in the P site^25^; or (3) stalling of the tRNA in the P site after translocation and/or EF-G dissociation^25,31-34^. Crystal structures of anticodon-stem-loops (ASLs) of +1FS-prone tRNAs in the A site^29,35-37^, formed in the absence of elongation factors, argue against the mechanism of +1FS during decoding, showing that steric hindrance in the decoding center prevents tRNA from slippage. Yet, the dynamics of the ribosome allow sampling of different structures, which may evade crystallization (e.g. refs^38,39^). Thus, the possibility of rearrangements of a frameshifting complex at all three elongation stages remain to be explored. To distinguish among these three possible mechanisms, it is necessary to capture ribosomal translocation complexes that are formed with full-length aminoacyl-tRNAs and EF-G on a +1FS-prone mRNA.

To visualize the structural mechanism of +1FS, we determined cryo-EM structures of 70S complexes with full-length native *E. coli* tRNA^Pro^(UGG) and EF-G, and compared the structures containing a non-frameshifting “control” mRNA with those containing a +1FS-prone mRNA. Unlike the ASLs of +1FS tRNAs that were used in previous studies^29,35-37^ and contained an extra nucleotide next to the anticodon^40^, native *E. coli* tRNA^Pro^(UGG) has a canonical anticodon loop. We first formed two pre-translocation 70S complexes, containing a non-frameshifting mRNA codon: C^1^CA-A^4^, or the frameshifting codon: C^1^CC-A^4^, in the A site. Each complex was prepared with fMet-tRNA^fMet^ in the P site and Pro-tRNA^Pro^(UGG) was delivered by EF-Tu•GTP to the A site. To capture EF-G-catalyzed translocation states, we then added EF-G with the non-hydrolyzable GTP analog GDPCP (5’-guanosyl-β,γ-methylene-triphosphate) to each pre-translocation complex and performed single-particle cryo-EM analyses (Methods). We used maximum-likelihood classification of cryo-EM data, which allows separation of numerous functional and conformational states within a single sample^41-43^. Our data classification revealed three elongation states in each complex (Figures S1 and S2): (1) pre-translocation structures with tRNA^Pro^ in the A site (I: non-frameshifting, and I-FS: frame<u>s</u>hifting); (2) “mid-translocation” EF-G-bound structure, with tRNA^Pro^ near the P site (II and II-FS); and (3) nearly fully translocated EF-G-bound state with tRNA^Pro^ in the P site (III and III-FS). Comparison of the non-frameshifting and frameshifting structures reveals that the ribosome is pre-disposed for +1FS before translocation, and that frameshifting is accomplished by the mid-translocation stage of EF-G-catalyzed translocation.

## RESULTS AND DISCUSSION

### Pre-translocation frameshifting structure I-FS adopts an open 30S conformation

Decoding of mRNA occurs on the non-rotated ribosome, in which peptidyl-tRNA occupies the P site and the aminoacyl-tRNA is delivered by EF-Tu to the A site^44-46^. Universally conserved 16S ribosomal RNA nucleotides of the decoding center G530, A1492 and A1493 (*E. coli* numbering) interact with the codon-anticodon helix, resulting in the closure of the 30S domain^47^, which stabilizes the cognate aminoacyl-tRNA during decoding^48,49^. Peptidyl transfer results in a deacylated tRNA in the P site and the peptidyl-tRNA in the A site, preparing the ribosome for translocation^6,46^. Thus, the closure of the 30S domain is a signature of canonical decoding at the A site.

We formed pre-translocation complexes by delivering Pro-tRNA^Pro^ with EF-Tu•GTP to the ribosomal A site containing the non-frameshifting CCA-A or frameshifting CCC-A motifs (Figure 1). Cryo-EM data classification reveals differences between the non-frameshifting and frameshifting complexes (Figure 1). While particle populations are similar (∼11% and ∼12%, respectively), consistent with comparable efficiency of decoding of both mRNA sequences^21^, the resulting cryo-EM maps report different conformations of the 30S subunit. The non-frameshifting Structure I features a canonical closed 30S subunit with G530, A1492 and A1493 in the ON state^48^, interacting with the backbone of the cognate codon-anticodon helix (Figures 1B-C). G530 contacts A1492, resulting in a latched decoding center. This conformation is nearly identical to that in other cognate 70S complexes^7,50,51^. By contrast, the frameshifting Structure I-FS with the U34-C3 wobble pair features an open 30S conformation (Figures 1E-F), in which the shoulder domain is shifted away from the body domain. This open conformation resembles transient intermediates of decoding captured by cryo-EM^39,48,49^. Here, G530 (at the shoulder) is retracted by ∼2 Å from the ON position, shifting away from A1492 (at the body) and from the backbone of G35 of tRNA^Pro^ (Figure 1F). Thus, the decoding-center triad is disrupted and provides weaker support for the codon-anticodon helix than in the non-frameshifting structure (Figure 1C). Structure I-FS therefore reveals that although the codon-anticodon helix is in the normal 0-frame (Figure 1E), the U34-C3 wobble pairing shifts the 30S dynamics equilibrium toward the open 30S conformation.

**Figure 1.**
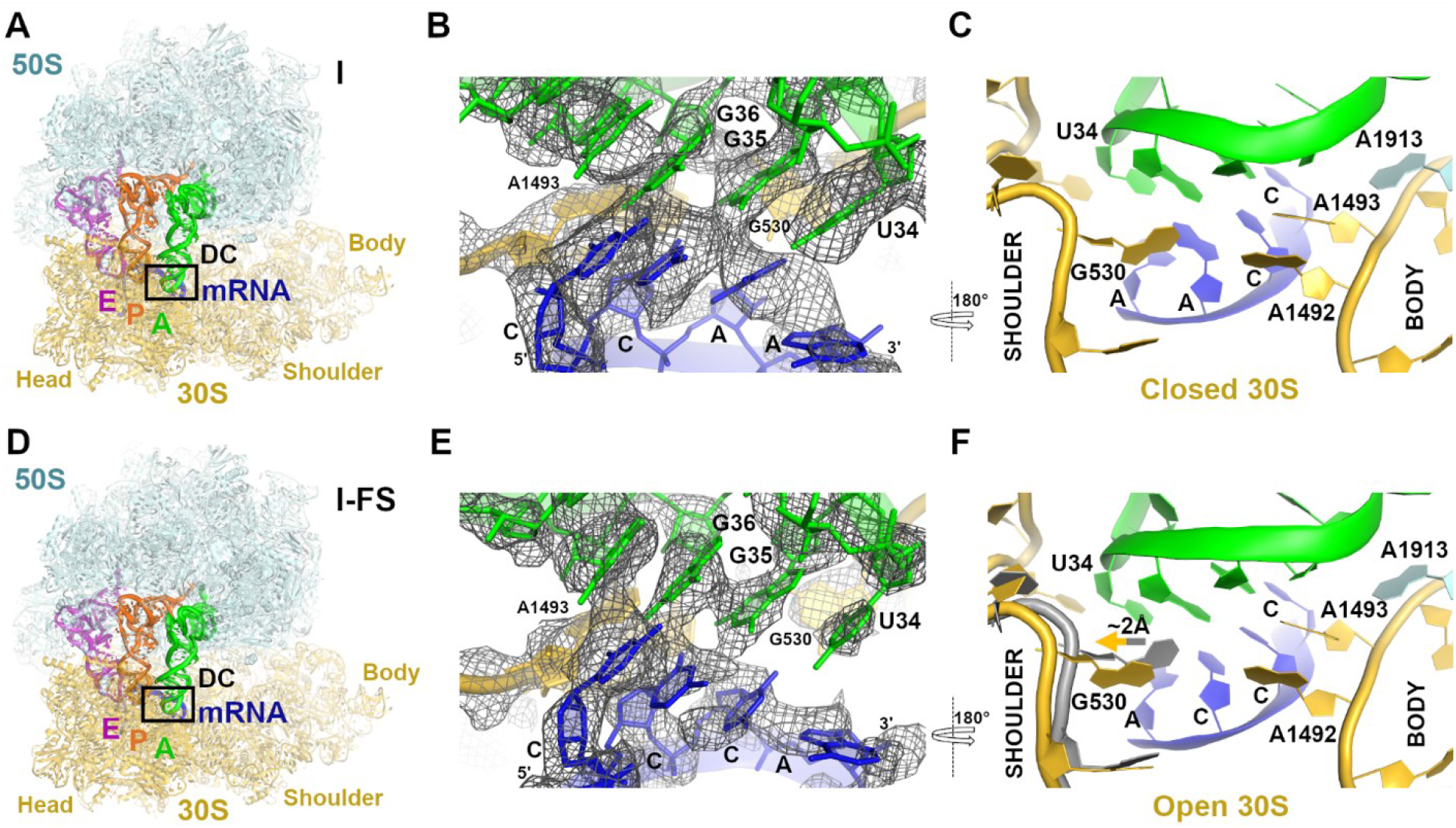
Cryo-EM structures of pre-translocation 70S formed with fMet-tRNA^fMet^ (P site) and Pro-tRNA^Pro^ (A site). (**A**) Overall view of the 70S structure with non-frameshifting mRNA (CCA-A; Structure I). Weaker density in the E site than in the A and P sites suggests partial occupancy of E-tRNA (Methods). (**B**) Cryo-EM density (gray mesh) for codon-anticodon interaction between non-frameshifting mRNA and tRNA^Pro^ in the A site of Structure I. The view approximately corresponds to the boxed decoding center region (DC) in panel A. The map was sharpened with a B-factor of -80 Å^2^ and is shown at 2.5 σ. (**C**) Decoding center nucleotides G530 (in the shoulder region) and A1492-A1493 (in the body region) stabilize the codon-anticodon helix in Structure I. (**D**) Overall view of the 70S structure with the slippery mRNA (CCC-A; Structure I-FS). Weaker density in the E site than in the A and P sites suggests partial occupancy of E-tRNA (see Methods). (**E**) Cryo-EM density for codon-anticodon interaction between the slippery mRNA codon and tRNA^Pro^ in Structure I-FS. The map was sharpened with a B-factor of -80 Å^2^ and is shown at 2.5 σ. (**F**) Partially open conformation of the 30S subunit due to the shifted G530 (in the shoulder region) in Structure I-FS relative to that in Structure I (16S shown in gray). Structural alignment was obtained by superposition of 16S ribosomal RNAs (rRNAs).

### mRNA frame is shifted in the EF-G-bound structures II-FS and III-FS

After peptidyl transfer, the pre-translocation 70S undergoes a thermally driven spontaneous rotation of the 30S subunit relative to the 50S subunit, allowing the tRNA acceptor arms to shift within the 50S subunit and adopt the hybrid A/P and P/E states^52^. EF-G•GTP binds to the rotated state^5,6,45,53,54^. Spontaneous reverse rotation of the 30S subunit in the presence of EF-G causes synchronous translocation of tRNA ASLs and mRNA codons within the 30S subunit, resulting in P/P and E/E states upon completion of the rotation^55^. Previous structures of 70S•2tRNA•EF-G complexes captured 30S in rotation states that ranged from ∼10 degrees to 0 degrees^53,56-58^, revealing early (rotated) and late (non-rotated) stages of translocation. They show that domain IV of EF-G binds next to the translocating peptidyl-tRNA and sterically hinders its return to the A site on the 30S subunit upon reverse subunit rotation^5,54,59^.

Our cryo-EM structures reveal two predominant translocation states with EF-G•GDPCP: the partially rotated state (∼5°) and the nearly non-rotated state (∼1°; relative to the non-rotated pre-translocation structure I) (Figures 2 and 3). The non-frameshifting structures II and III closely resemble previously described mid-translocated^57,58^ (Figures 2A-C) and post-translocated^56^ structures (Figures 3A-B) formed with antibiotics. In the partially rotated state, the head of the 30S subunit is swiveled by ∼16°, so the 30S beak is closer to the 50S subunit (Figure 2A). The head swivel is coupled with tRNA ASL and mRNA translocation on the small subunit, allowing gradual translocation first relative to the 30S body then 30S head^60^. In the head-swiveled Structure II, dipeptidyl fMet-Pro-tRNA^Pro^ is between the A and P sites of the 30S subunit (Figure 2B). Specifically, the anticodon nucleotide U34 is ∼4 Å away from the P site of the body domain. Yet, the anticodon remains near the A site of the head domain due to the movement of the head in the direction of translocation. The acceptor arm is in the P site of the 50S subunit. Thus, the tRNA conformation is similar to the previously described chimeric ap/P conformation^57^ (denoting the anticodon at the A site of the 30S head and near the P site of the 30S body (ap), and the acceptor arm in the P site of the 50S subunit (P)).

**Figure 2.**
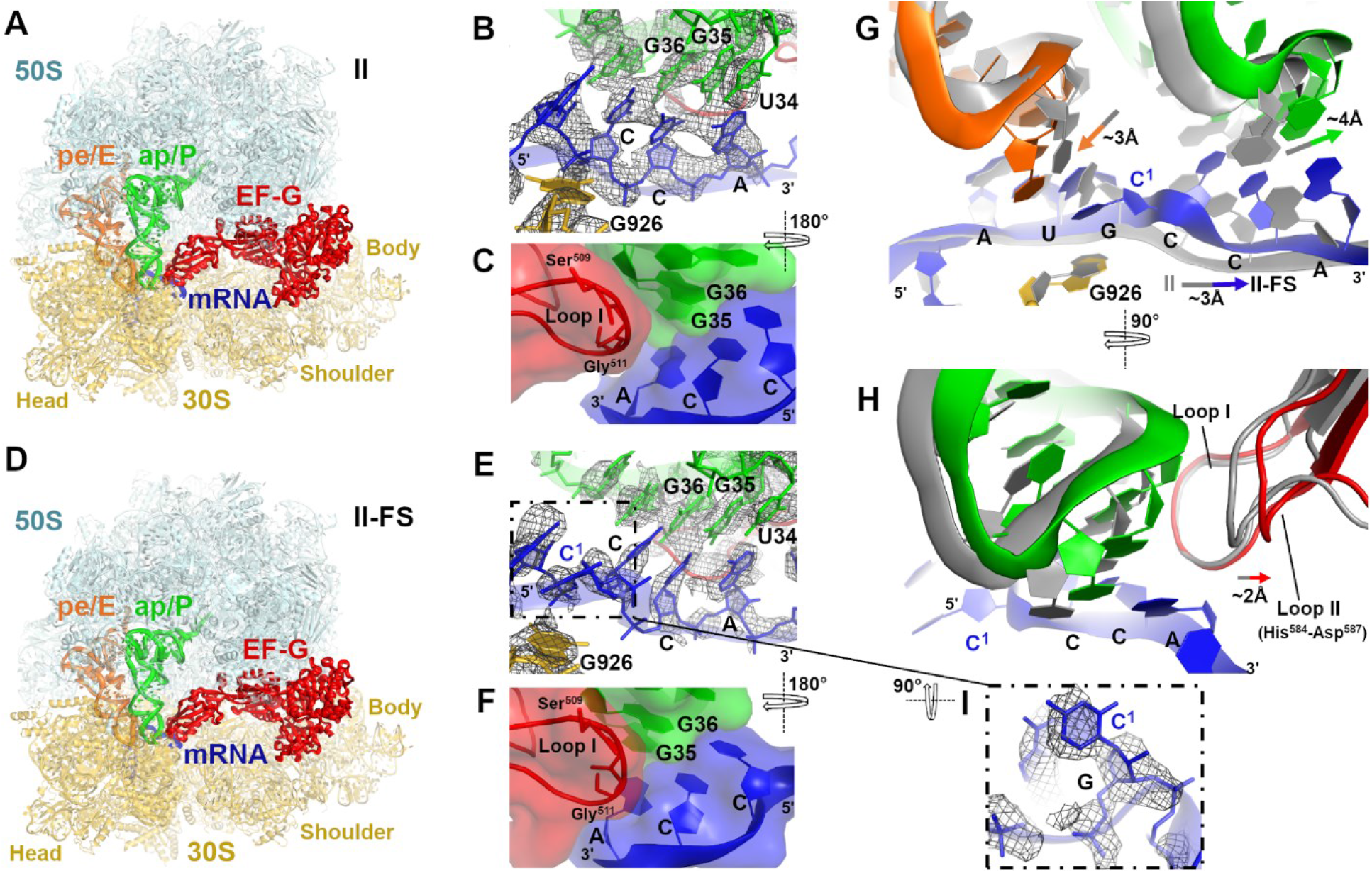
Cryo-EM structures of mid-translocation states formed with EF-G•GDPCP. (**A**) Overall view of mid-translocation Structure II with the non-frameshifting mRNA. (**B**) Cryo-EM density (mesh) of the non-frameshifting tRNA^Pro^ and mRNA codon near the P site. The map is sharpened by applying the B-factor of-80 Å^2^ and is shown with 2.5 σ. (**C**) Interaction of the EF-G loop I (Ser509-Gly51, red) with the codon-anticodon helix (space-filling surface and cartoon representation). (**D**) Overall view of mid-translocation Structure II-FS with the frameshifting mRNA. (**E**) Cryo-EM density of the frameshifting tRNA^Pro^ and mRNA codon near the P site. The map is sharpened by applying the B-factor of -80 Å^2^ and is shown with 2.5 σ. Note the unpaired and bulged C^1^ nucleotide in the mRNA, also shown in panel I. (**F**) Interaction of the EF-G loop I (Ser509-Gly510) with the codon-anticodon helix of the frameshifting mRNA (compare to panel **C**). (**G**) Differences in positions of tRNA^Pro^ (green) and tRNA^fMet^ (orange) in the frameshifting structure II-FS relative to those in the non-frameshifted structure II (gray). (**H**) Adjustment of loop II of EF-G (red) to accommodate the shifted position of tRNA^Pro^ (green) in the frameshifting structure II-FS relative to those in structure II (gray). Structural alignments were performed by superposition of 16S rRNAs. (**I**) Close-up view of cryo-EM density for bulged C1 in Structure II-FS (also shown in panel E).

**Figure 3.**
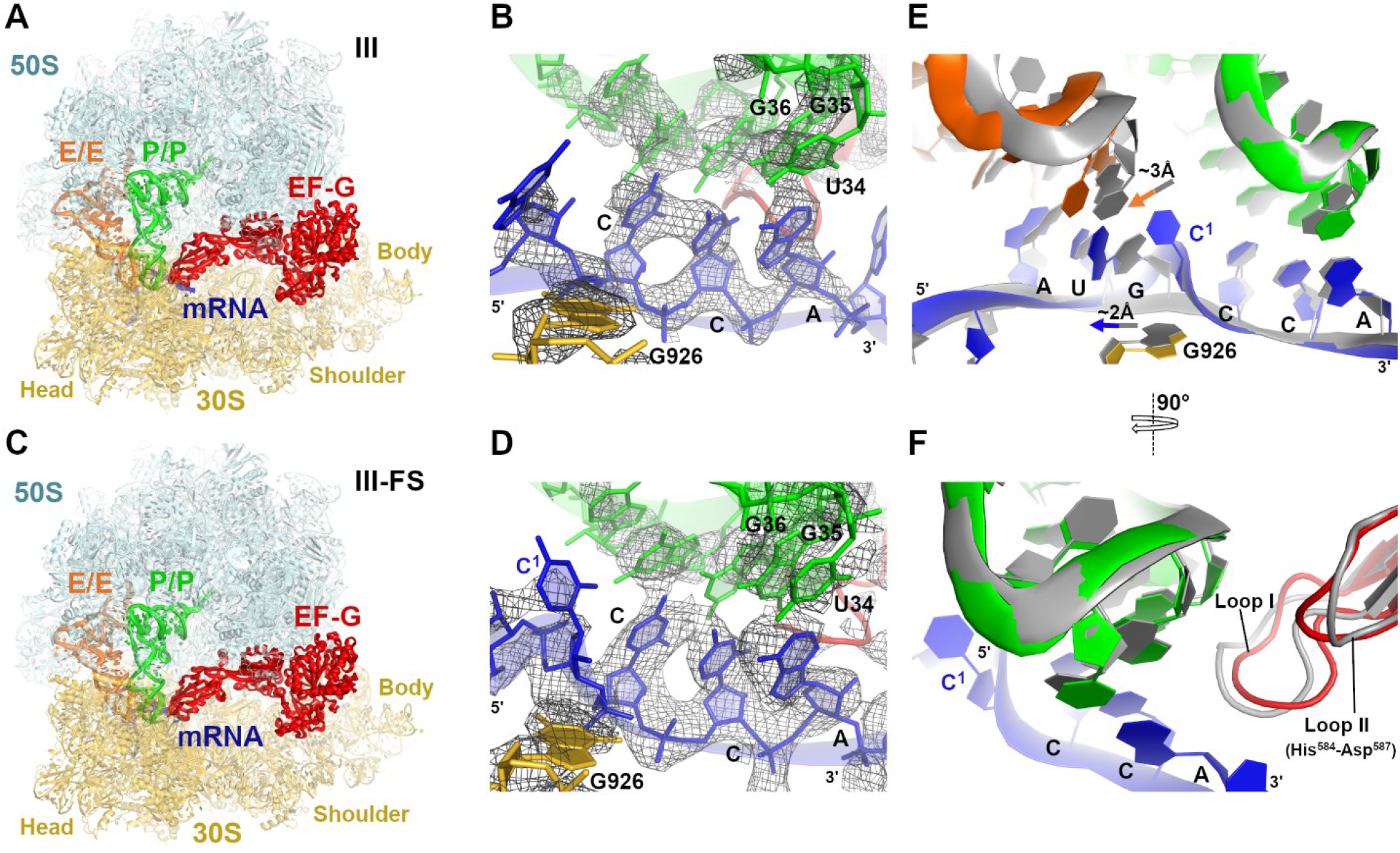
Cryo-EM structures of near post-translocation states formed with EF-G•GDPCP. (**A**) Overall view of the near-post-translocation Structure III with the non-frameshifting mRNA. (**B**) Cryo-EM density (mesh) of the non-frameshifted tRNA^Pro^ and mRNA codon at the P site. The map was sharpened by applying the B-factor of -80 Å^2^ and is shown at 2.5 σ. (**C**) Overall view of the near post-translocation Structure III-FS with the frameshifting mRNA. (**D**) Cryo-EM density (mesh) of the frameshifted tRNA^Pro^ and mRNA codon at the P site. The map was sharpened by applying the B-factor of -80 Å^2^ and is shown at 2.5 σ. (**E**) Comparison of mRNA and tRNA positions in the nearly translocated frameshifted (colored, III-FS) and non-frameshifted (gray, III) complexes. (**F**) Positions of loop II of EF-G (red) and tRNA^Pro^ (green) in the Structures III-FS and III (gray). Structural alignments were performed by superposition of 16S rRNAs.

The nearly non-rotated Structure III features a small head swivel (∼1°) and dipeptidyl-tRNA in the P site (Figures 3A-B and S3), resembling the non-rotated post-translocation ribosome^56^. Both the dipeptidyl-tRNA^Pro^ and the deacylated tRNA^fMet^ are base-paired with their respective mRNA codons in the P and E sites, respectively. In both structures II and III, domain IV of EF-G interacts with the anticodon stem-loop of the dipeptidyl-tRNA and the proline CCA codon (Figures 2B-C and 3B), consistent with the role of EF-G in stabilizing the codon-anticodon helix during translocation^57,61^ and after arrival of the codon-anticodon helix at the P site^56^.

In contrast to the non-frameshifting complex, EF-G•GDPCP mediates frameshifting on the frameshifting mRNA. In the mid-translocated Structure II-FS, the dipeptidyl-tRNA^Pro^ (Figure S3B) pairs with the mRNA in the +1-frame (C^2^CA^4^) between the A and P sites of the 30S subunit (Figures 2D-F). Here, clearly resolved density demonstrates base-pairing of U34 of tRNA^Pro^ with A4 of the mRNA (Figure 2E). The neighboring deacylated tRNA^fMet^ is bound to the AUG codon near the E site. Thus, +1 frameshifting results in a bulged mRNA nucleotide C1 between the E and P sites (Figures 2E, 2G, 2I). C1 is sandwiched between the guanosine of the AUG codon and G926 of 16S rRNA. This stabilization allows mRNA compaction and accommodation of four mRNA nucleotides in the E-site, which normally accommodates three nucleotides^7^. Due to frameshifting, tRNA^fMet^ and tRNA^Pro^ are shifted away from each other; they are moved by 3 Å and 4 Å from their positions in the non-frameshifting Structure II, respectively (Figure 2G). The shift of tRNA^Pro^ is compensated by the shift of loop II of EF-G, whereas the rest of domain IV is placed similarly to that in the non-frameshifting complex (Figure 2H).

Previous crystallographic work suggested that the 16S rRNA nucleotides C1397 and A1503, which flank the A and E sites, respectively, prevent mRNA slippage by interacting with the bases of translocating mRNA^58,61,62^. These two nucleotides are part of the central region of the 30S head that is stabilized by numerous interactions, such as the conserved 1399-1504 Watson-Crick base pair formed by nucleotides neighboring the “stoppers” C1397 and A1503. Our structures indicate that the positions and conformations of this head region, including C1397 and A1503, are nearly identical in the non-frameshifting Structure II as in the frameshifted Structure II-FS (Figure S4). Thus, the compact and frameshifted mRNA can be accommodated in the ribosomal mRNA tunnel during elongation without perturbing the conformations of the head nucleotides.

In the nearly translocated non-rotated Structure III-FS (Figure 3C), the frameshifted CCA codon and dipeptidyl-tRNA^Pro^ are in the P site, while C1 and the AUG codon with the deacylated tRNA^fMet^ are in the E site (Figure 3D). To accommodate C1 in the E site, the E-site AUG codon and the paired tRNA^fMet^ are shifted by up to 3 Å (Figure 3E). Weak C1 density suggests that C1 is detached from G926, which instead hydrogen-bonds with the phosphate group of the first nucleotide of the P-site codon (Figures 3D-E). The P-site codon and tRNA^Pro^ are positioned nearly identically to those in the non-frameshifting Structure III (Figure 3F). Thus, the frameshifted mRNA and peptidyl-tRNA are placed at the canonical P-site position at the end of the translocation trajectory, preparing the ribosome for the next elongation cycle on the new +1-frame of the mRNA.

### Mechanism of +1 frameshifting

Cryo-EM structures in this work provide the long-sought snapshots of +1 frameshifting (Figure 4). The use of the native *E. coli* tRNA and visualization of EF-G-bound structures distinguishes this work from previous structural studies that were based on +1-frameshift suppressor tRNAs with an expanded anticodon loop^29,34-37^ or frameshifting-like complexes with a single tRNA^34,63^. To obtain a complete elongation complex with two tRNAs required for translocation, that would be prone to +1FS, we used a frameshifting mRNA sequence C^1^CC-A^4^ and tRNA^Pro^ (UGG)^21^ in the A site. The frameshifting ribosome complex therefore contains a wobble U34-C3 pair upon binding of tRNA^Pro^ to the C^1^CC-A^4^ sequence (Structure I-FS). While the downstream A4 would have been a more favorable base-pairing partner for U34 of tRNA^Pro^, there is no frameshifting upon decoding. Thus, the +1FS-prone pre-translocation complex maintains the 0-frame anticodon-codon pairing resembling that in canonical elongation complexes^51^ and crystal structures with suppressor tRNAs^35-37^. However, unlike the non-frameshifting complex containing the U34-A3 base pair (Structure I) and unlike previous structures with suppressor tRNAs^35-37^ structure I-FS features an open 30S subunit, resembling transient decoding intermediates^48,49^. Here, G530 of 16S rRNA is shifted from its canonical position near the second base pair of the codon-anticodon helix^47^, thus possibly destabilizing the labile three-base-pair codon-anticodon helix. This structure appears pre-disposed for tRNA^Pro^ to slide from its near-cognate codon CCC to the cognate CCA codon in the +1-frame.

**Figure 4.**
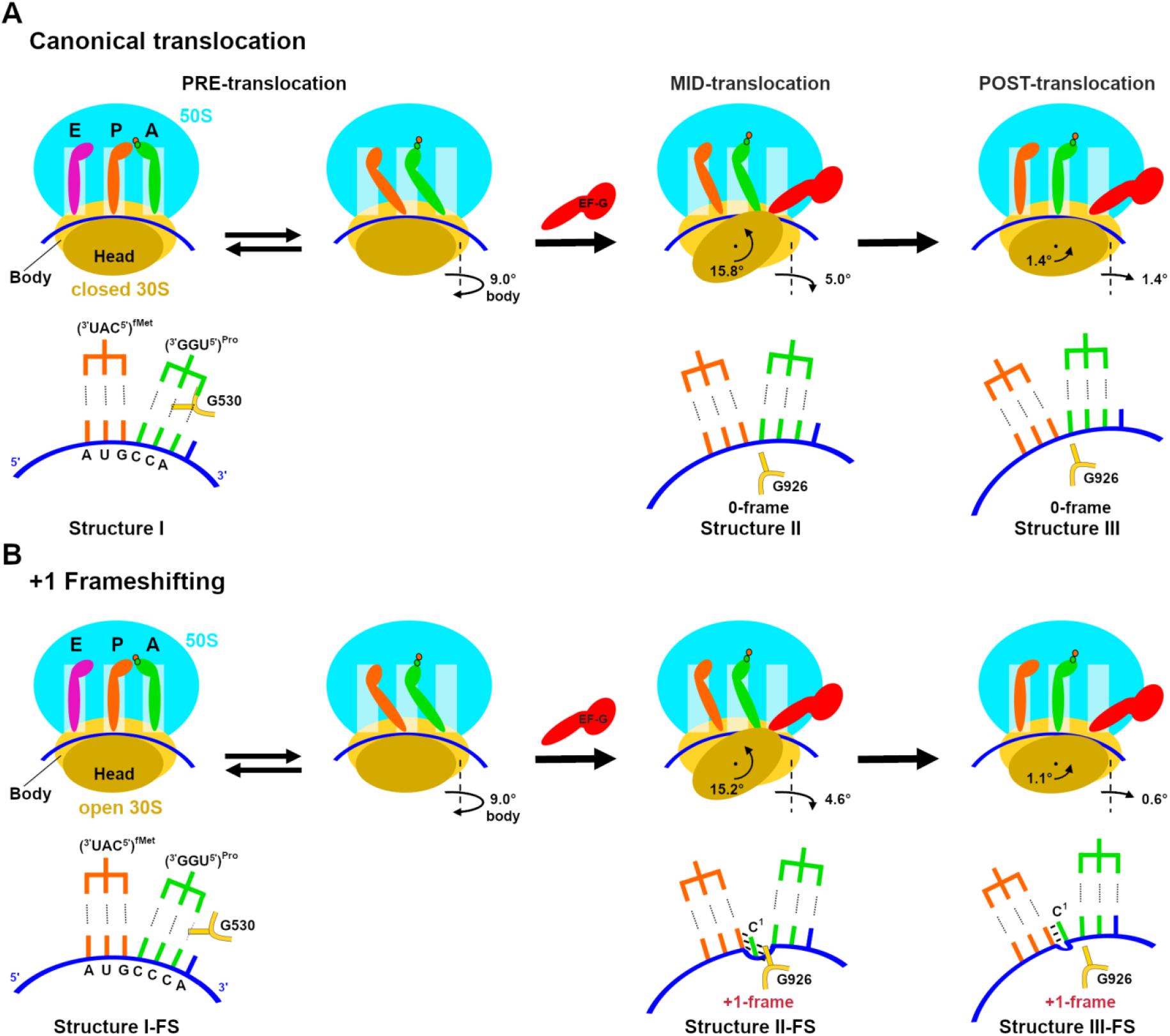
The mechanism of +1 frameshifting. (**A**) Schematic of canonical ribosomal translocation by EF-G and ribosome rearrangements. (**B**) Schematic of ribosomal translocation by EF-G resulting in +1 frameshifting. The second rows in panels A and B show local rearrangements of mRNA-tRNA and positions of the decoding-center nucleotide G530 and P-site nucleotide G926 of the 30S subunit.

Limited space in the A site, however, restricts the codon-anticodon dynamics and prevents slippage in this pre-translocation state. In contrast, the mid-translocation complex with EF-G and the highly swiveled 30S head features tRNA^Pro^ base-paired with the C^2^CA^4^ codon near the P site of the body and A site of the head (Structure II-FS). This indicates that the ribosome switches to the +1-frame when tRNA^Pro^ and mRNA move from the decoding center, and that frameshifting is accomplished by the intermediate of EF-G-catalyzed translocation, at which the tRNA is nearly translocated along the 30S body. The complex remains frameshifted till the completion of translocation when tRNA^Pro^ is in the P site relative to both the body and head due to the reverse head swivel (Structure III-FS). Our work therefore suggests a structural mechanism, in which non-canonical pairing of the pre-translocation complex sets the stage for frameshifting by opening the 30S subunit and promoting frameshifting during EF-G-catalyzed translocation.

Our observation of the destabilization of the pre-translocation complex, and the EF-G-bound frameshifting structures, is consistent with the high efficiency of +1 frameshifting on the CCC-A frameshifting codon motif shown *in vitro*^21^. Other frameshifting sequences exist, however, which contain fully complementary codon-anticodon interactions in the 0- and +1-frames, such as the CCC-C sequence decoded by tRNA^Pro^ (GGG)^25,64^. In these latter cases, the pre-translocation complex most likely samples a canonical closed 30S conformation, in which the codon-anticodon helix is stabilized by the decoding center (as in Structure I). This frame stabilization must at least in part account for the lower efficiency of frameshifting on such sequences^21,64^. Nevertheless, the low frequency with which +1 frameshifting occurs with such sequences indicates that the tRNA-mRNA interactions can be stochastically destabilized during translocation, when the small subunit, tRNAs, and mRNA rearrange. Indeed, recent 70S structures obtained without EF-G demonstrate that mRNA frame destabilization occurs upon 30S head swiveling. In a frameshift-like complex featuring a single tRNA and swiveled 30S head, the bulged nucleotide between the E and P site codons is stabilized by G926^34^, similarly to that in Structure II-FS. Furthermore, a recent crystal structure of a non-frameshifting complex with two tRNAs and swiveled 30S head revealed perturbation of the codon-anticodon interactions in the P site, despite full complementarity of the P-site tRNA with the 0-frame codon^62^. While tRNA-mRNA pairing is unstable during head swiveling, EF-G maintains the reading frame in non-frameshifting complexes by interacting with both the tRNA anticodon and mRNA codon along the translocation trajectory (Structures II and III). By contrast, in the frameshifting-prone complexes, EF-G fails to support the codon-anticodon interactions destabilized in the initial stages of translocation (such as CCC-A in this study) and allows slippage into the fully complementary sequences that can pair with tRNA in the +1-frame.

## METHODS

### Preparation of EF-G and ribosomal subunits

The gene encoding full-length *E. coli* EF-G (704 aa, C-terminally His_6_-tagged) cloned into pET24a+ plasmid (Novagen, kanamycin resistance vector) was transformed into an *E. coli* BLR/DE3 strain. Cells with the plasmid were cultured in Luria-Bertani (LB) medium with 50 µg mL^-1^ kanamycin at 37 °C until the OD_600_ reached 0.7-0.8. Expression of EF-G was induced by 1 mM IPTG (Gold Biotechnology Inc., USA), followed by cell growth for 9 hrs at 16 °C. The cells were harvested, washed and resuspended in buffer A (50 mM Tris pH=7.5, 50 mM NH_4_Cl, 10 mM MgCl_2_, 5% glycerol, 10 mM imidazole, 6 mM β-mercaptoethanol (βME), and a cocktail of protease inhibitors (complete Mini, EDTA-free protease inhibitor tablets, Sigma Aldrich, USA). The cells were disrupted with a microfluidizer (Microfluidics, USA), and the soluble fraction was collected by centrifugation at 18,000 rpm for 50 minutes and filtered through a 0.22 μm pore size sterile filter (CELLTREAT Scientific Products, USA).

EF-G was purified in three steps. The purity of the protein after each step was verified by 12 % SDS-PAGE stained with Coomassie Brilliant Blue R 250 (Sigma-Aldrich). First, affinity chromatography with Ni-NTA column (Nickel-nitrilotriacetic acid, 5 ml HisTrap, GE Healthcare) was performed using FPLC (Äkta explorer, GE Healthcare). The soluble fraction of cell lysates was loaded onto the column equilibrated with buffer A and washed with the same buffer. EF-G was eluted with a linear gradient of buffer B (buffer A with 0.25 M imidazole). Fractions containing EF-G were pooled and dialyzed against buffer C (50mM Tris pH=7.5, 100 mM KCl, 10 mM MgCl_2_, 0.5 mM EDTA, 6 mM βME, and the cocktail of protease inhibitors). The protein then was purified by ion-exchange chromatography through a HiPrep FF Q-column (20 mL, GE Healthcare; FPLC). After the column was equilibrated and washed with Buffer C, the protein was loaded in Buffer C and eluted with a linear gradient of Buffer D (Buffer C with 0.7 M KCl). Finally, the protein was dialyzed against 50 mM Tris pH=7.5, 100 mM KCl, 10 mM MgCl_2_, 0.5 mM EDTA, 6 mM βME, and purified using size-exclusion chromatography (Hiload 16/600 Superdex 200pg column, GE Healthcare). The fractions of the protein were pooled, buffer exchanged (25 mM Tris pH=7.5, 100 mM NH_4_Cl, 10 mM MgCl_2_, 0.5 mM EDTA, 6 mM βME, and 5% glycerol) and concentrated with an ultrafiltration unit using a 10-kDa cutoff membrane (Millipore). The concentrated protein was flash-frozen in liquid nitrogen and stored at -80 ^°^C.

70S ribosomes were prepared from *E. coli* (MRE600) as described^65^, and stored in the ribosome-storage buffer (20 mM Tris-HCl (pH 7.0), 100 mM NH_4_Cl, 12.5 mM MgCl_2_, 0.5 mM EDTA, 6 mM βME) at -80°C. Ribosomal 30S and 50S subunits were purified using a sucrose gradient (10-35%) in a ribosome-dissociation buffer (20 mM Tris-HCl (pH 7.0), 500 mM NH_4_Cl, 1.5 mM MgCl_2_, 0.5mM EDTA, and 6 mM βME). The fractions containing 30S and 50S subunits were collected separately, concentrated and stored in the ribosome-storage buffer at -80°C.

### Preparation of charged tRNAs, and mRNA sequences

*E. coli* tRNA^fMet^ was purchased from Chemical Block and aminoacylated as described^66^.

Native *E. coli* tRNA^Pro^(UGG) (*proM* tRNA) was over-expressed in *E. coli* from an IPTG-inducible *proM* gene carried by pKK223-3. Total tRNA was isolated using differential centrifugation^67^ and *proM* tRNA was isolated using a complementary biotinylated oligonucleotide attached to streptavidin-sepharose^68^ yielding approximately 40 nmoles *proM* tRNA from 1 liter of culture. *E. coli* tRNA^Pro^ (UGG) (10 µM) was aminoacylated in the charging buffer (50 mM Hepes pH 7.5, 50 mM KCl, 10 mM MgCl_2_, 10 mM DTT) in the presence of 40 µM L-proline, 2 µM prolyl-tRNA synthetase (ProRS), 0.625 mM ATP and 15 µM elongation factor EF-Tu (purified as described^48^). The mixture was incubated for 10 minutes at 37°C. To stabilize the charged Pro-tRNA^Pro^ and form the ternary complex for the elongation reaction 0.25 mM GTP was added to the mixture. The mixture was incubated for 3 minutes at 37°C.

mRNAs containing the Shine-Dalgarno sequence and a linker to place the AUG codon in the P site were synthesized by IDT. The frameshifting mRNA contains the sequence 5’-GGC AAG GAG GUA AAA <u>AUG</u> **CCC** AGU UCU AAA AAA AAA AAA, and the non-frameshifting mRNA contains the sequence 5’-GGC AAG GAG GUA AAA <u>AUG</u> **CCA** AGU UCU AAA AAA AAA AAA.

### Preparation of 70S translocation complexes with EF-G•GDPCP

70S•mRNA•fMet-tRNA^fMet^•Pro-tRNA^Pro^(UGG)•EF-G•GDPCP complexes were prepared as follows, separately for the slippery and non-slippery mRNAs. In each, 0.33 µM 30S subunits (all concentrations specified for the final solution) were pre-activated at 42°C for 5 minutes in the ribosome-reconstitution buffer (20 mM HEPES pH 7.5, 120 mM NH_4_Cl, 20mM MgCl_2_, 2 mM spermidine, 0.05 mM spermine, 6 mM βME). These activated 30S subunits were added with0.33 µM 50S subunits with 1.33 µM mRNA and incubated for 10 minutes at 37°C. Subsequently, 0.33 µM fMet-tRNA^fMet^ was added and the solution was incubated for 3 minutes at 37°C, to form the 70S complex with the P-site tRNA.

Pro-tRNA^Pro^ (UGG) (0.33 µM), EF-Tu (0.5 µM), and GTP (8.3 µM) were added to the solution and incubated for 10 minutes at 37°C to form the A-site bound 70S complex. Next, EF-G (5.3 µM) and GDPCP (0.66 mM) were added and incubated for 5 minutes at 37°C, then cooled down to room temperature, resulting in 70S translocation complexes with EF-G•GDPCP.

### Cryo-EM and image processing

QUANTIFOIL R 2/1 grids with the 2-nm carbon layer (Cu 200, Quantifoil Micro Tools) were glow discharged with 25 mA with negative polarity for 60 s in a PELCO easiGlow glow discharge unit. Each complex (2.5 μL) was separately applied to the grids. Grids were blotted at blotting force 9 for 4 s at 5°C, 95% humidity, and plunged into liquid ethane using a Vitrobot MK4 (FEI). Grids were stored in liquid nitrogen.

For the frameshifting 70S•mRNA(CCC-A)•fMet-tRNA^fMet^•Pro-tRNA^Pro^(UGG)•EF-G•GDPCP translocation complex, a dataset of 164,504 particles was collected as follows. A total of 2,591 movies were collected on Titan Krios (FEI) microscope (operating at 300 kV) equipped with K2 Summit camera system (Gatan), with −0.8 to -2.0 μm defocus. Multi-shot data collection was performed by recording four exposures per hole, using SerialEM^69^ with a beam-image shift, as described^70^. Coma-free alignment was performed using a built-in function. ‘Coma vs. Image Shift’ from the Calibration menu was used for dynamic beam-tilt compensation, based on image shifts for each exposure. Multi-shot configuration was selected from ‘Multiple Record Setup Dialog’ to dynamically adjust the beam tilt. Backlash-corrected compensation was applied to each stage movement at the target stage position to reduce mechanical stage drift. Each exposure was acquired with continuous frame streaming at 36 frames per 7.2 s, yielding a total dose of 47.5 e^-^/Å^2^. The dose rate was 7.39 e^-^/upix/s at the camera. The nominal magnification was 130,000 and the calibrated super-resolution pixel size at the specimen level was 0.525 Å. The movies were motion-corrected and frame averages were calculated using all 36 frames within each movie after multiplying by the corresponding gain reference in IMOD^71^. During motion-correction in IMOD the movies were binned to pixel size 1.05 Å (termed unbinned or 1×binned). cisTEM^72^ was used to determine defocus values for each resulting frame average and for particle picking. All movies were used for further analysis after inspection of the averages and the power spectra computed by CTFFIND4 within cisTEM. The stack and particle parameter files were assembled in cisTEM with the binnings of 1×, 2× and 4× (box size of 400 for unbinned stack). Data classification is summarized in Figure S2. FREALIGNX was used for all steps of particle alignment, refinement and final reconstruction steps and FREALIGN v9.11 was used for 3D classification steps^73^. Conversion of parameter file from FREALIGNX to FREALIGN for classification was performed by removing a column twelve which contains phase shift information (not applicable as no phase plate was used) and adding an absolute magnification value. Reverse conversion from FREALIGN to FREALIGNX for refinement was performed automatically by FREALIGNX. The 4x-binned image stack (164,504 particles) was initially aligned to a ribosome reference (PDB 5U9F)^74^ using 5 cycles of mode 3 (global search) alignment including data in the resolution range from 300 Å to 30 Å until the convergence of the average score. Subsequently, the 4x binned stack was aligned against the common reference resulting from the previous step, using mode 1 (refine) in the resolution range 300-18 Å (3 cycles of mode 1). In the following steps, the 4x binned stack was replaced by the 2x binned image stack, which was successively aligned against the common reference using mode 1 (refine), including gradually increasing resolution limits (5 cycles per each resolution limit; 18-12-10-8 Å) up to 8 Å. 3D density reconstruction was obtained using 60% of particles with highest scores. Subsequently, the refined parameters were used for classification of the 2x binned stack into 16 classes in 50 cycles using the resolution range of 300-8 Å. This classification revealed 11 high-resolution classes, 3 low-resolution (junk) classes, and 2 classes representing only the 50S subunit (Figure S2A). The particles assigned to the high-resolution 70S classes were extracted from the 2x binned stack (with > 50% occupancy and scores > 0) using merge_classes.exe (part of the FREALIGN distribution), resulting in a stack containing 109,094 particles. Classification of this stack was performed for 50 cycles using a focused spherical mask between the A and P sites (30 Å radius, as implemented in FREALIGN). This sub-classification into 8 classes yielded one high-resolution class, which contained both tRNAs and EF-G; and one high-resolution class, which contained 3 tRNAs (Structure I-FS). The map corresponding to an EF-G-bound translocation state had heterogeneous 30S features corresponding to a mixture of two states (with a highly swiveled and less-swiveled head conformations). The particles assigned to the high-resolution class with both tRNAs and EF-G were extracted from the 2x binned stack (with > 50% occupancy and scores > 0) using merge_classes.exe (part of the FREALIGN distribution), resulting in a stack containing 15,088 particles. Classification of this stack was performed for 50 cycles using a 3D mask designed around the head of 30S subunit. This sub-classification into 2 classes yielded 2 high-resolution classes, which contained both tRNAs and EF-G but differed in 30S head rotation (Structure II-FS and III-FS). Using subsequent sub-classification of each class into more classes did not yield additional structures. For the classes of interest (Structure I-FS, 12,108 particles; Structure II-FS, 9,059 particles; Structure III-FS, 6,029 particles), particles with > 50% occupancy and scores > 0 were extracted from the 2x binned stack. Refinement to 6 Å resolution using mode 1 (5 cycles) of the respective 1x binned stack using 95% of particles with highest scores resulted in ∼3.2 Å (Structure I-FS), ∼3.2 Å (Structure II-FS) and ∼3.3 Å (Structure III-FS) maps (FSC=0.143).

For the non-frameshifting 70S•mRNA(CCA-A)•fMet-tRNA^fMet^•Pro-tRNA^Pro^ (UGG)•EF-G•GDPCP translocation complex, a dataset of 1,041 movies containing 62,716 particles was collected and processed the same way as that for the frameshifting complex. All movies were used for further analysis after inspection of the averages and the power spectra computed by CTFFIND4 within cisTEM. The stack and particle parameter files were assembled in cisTEM with the binnings of 1×, 2× and 4× (box size of 400 for a unbinned stack). Data classification is summarized in Figure S1. FREALIGNX was used for all steps of particle alignment, refinement and final reconstruction steps and FREALIGN v9.11 was used for 3D classification steps ^73^ as described above. The 4x-binned image stack (62,716 particles) was initially aligned to a ribosome reference (PDB 5U9F)^74^ using 5 cycles of mode 3 (global search) alignment including data in the resolution range from 300 Å to 30 Å until the convergence of the average score. Subsequently, the 4x binned stack was aligned against the common reference resulting from the previous step, using mode 1 (refine) in the resolution range 300-18 Å (3 cycles of mode 1). In the following steps, the 4x binned stack was replaced by the 2x binned image stack, which was successively aligned against the common reference using mode 1 (refine), including gradually increasing resolution limits (5 cycles per each resolution limit; 18-12-10-8 Å) up to 8 Å. 3D density reconstruction was obtained using 60% of particles with highest scores. The refined parameters were used for classification of the 2x binned stack into 8 classes in 50 cycles using the resolution range of 300-8 Å. This classification revealed six high-resolution classes, one low-resolution (junk) class, and one class representing only 50S subunit (Figure S1A). The particles assigned to the high-resolution 70S classes were extracted from the 2x binned stack (with > 50% occupancy and scores > 0) using merge_classes.exe (part of the FREALIGN distribution), resulting in a stack containing 41,382 particles. Classification of this stack was performed for 50 cycles using a focused spherical mask between the A and P sites (30 Å radius, as implemented in FREALIGN). This sub-classification into eight classes yielded two high-resolution classes, which contained both tRNAs and EF-G (Structure II and III); and one high-resolution class, which contained 3 tRNAs (Structure I). For the classes of interest (Structure I, 4,263 particles; Structure II, 3,179 particles; Structure III, 4,612 particles), particles with > 50% occupancy and scores > 0 were extracted from the 2x binned stack. Refinement to 6 Å resolution using mode 1 (5 cycles) of the respective 1x binned stack using 95% of particles with highest scores resulted in ∼3.4 Å (Structure I), ∼3.5 Å (Structure II) and ∼3.4 Å (Structure III) maps (FSC=0.143). In both Structures I and I-FS, E-tRNA density is weak, indicating partial E-site occupancy. This is similar to our previous observations^39^, where additional classification resulted in maps with the vacant and tRNA-bound E-site, however no other differences (i.e. in the occupancy of other sites, or ribosome conformations) were observed. To account for partial density, we have modeled E-site tRNA, as described in^39^.

The maps (Structure I-FS, II-FS, III-FS, I, II and III) were filtered for structure refinements, by blocres and blocfilt from the Bsoft package^75^. Briefly, a mask was created for each map by low-pass filtering the map to 30 Å in Bsoft, then binarizing, expanding by 3 pixels and applying a 3-pixel Gaussian edge in EMAN2^76^. Blocres was run with a box size of 20 pixels for all maps. In each case, the resolution criterion was FSC with cutoff of 0.143. The output of blocres was used to filter maps according to local resolution using blocfilt. The optimal balance between high-resolution and lower-resolution regions is achieved for blocfilt maps filtered with a constant B-factor of -80 Å^2^ in bfactor.exe (part of the FREALIGN distribution). These B-factor sharpened maps were used for model building and structure refinements. Other B-factors were also used to interpret high-resolution details in the ribosome core regions. Fourier shell correlation (FSC) curves were calculated by FREALIGN for even and odd particle half-sets.

### Model building and refinement

Reported cryo-EM structure of *E. coli* 70S•fMet-tRNA^Met^•Phe-tRNA^Phe^•EF-Tu•GDPCP complex (PDB 5UYM)^48^, excluding EF-Tu and tRNAs, was used as a starting model for structure refinement. The structure of EF-G from PDB 4V7D^53^ was used as a starting model, and switch regions were generated by homology modeling from PDB 4V9P^77^. The structure of tRNA^Pro^ (UGG) was created by homology modeling (according to tRNA^Pro^ (UGG) sequence) using ribosome-bound tRNA^Pro^ (CGG) (PDB 6ENJ)^78^.

Initial protein and ribosome domain fitting into cryo-EM maps was performed using Chimera^79^, followed by manual modeling using PyMOL^80^. The linkers between the domains and parts of the domains that were not well defined in the cryo-EM maps (e.g. loops of EF-G) were not modeled.

All structures were refined by real-space simulated-annealing refinement using atomic electron scattering factors in RSRef^81,82^ as described^83^. Secondary-structure restraints, comprising hydrogen-bonding restraints for ribosomal proteins and base-pairing restraints for RNA molecules, were employed as described^84^. Refinement parameters, such as the relative weighting of stereochemical restraints and experimental energy term, were optimized to produce the stereochemically optimal models that closely agree with the corresponding maps. In the final stage, the structures were refined using phenix.real_space_refine^85^, followed by a round of refinement in RSRef applying harmonic restraints to preserve protein backbone geometry. The refined structural models closely agree with the corresponding maps, as indicated by low real-space R-factors and high correlation coefficients (Table S1). The resulting models have excellent stereochemical parameters, characterized by low deviation from ideal bond lengths and angles, low number of protein-backbone outliers and other robust structure-quality statistics, as shown in Table S1. Structure quality was validated using MolProbity^86^.

Structure superpositions and distance calculations were performed in PyMOL. To calculate the degree of the 30S body rotation or head rotation (swivel) between two 70S structures, the 23S rRNAs or 16S rRNAs of the 30S body were aligned using PyMOL, and the angle was measured in Chimera. These degrees of rotation (30S body/subunit rotation and 30S head rotation) for Structures II, III, II-FS and III-FS are reported relative to the classical non-rotated Structures I and I-FS, respectively. Figures were prepared in PyMOL and Chimera^79,80^. PDB coordinates were deposited in RCSB and cryo-EM maps were deposited in EMDB. Structure I: PDBID:7K50 EMDB:EMD-22669; Structure II: PDBID:7K51 EMDB:EMD-22670; Structure III: PDBID:7K52 EMDB:EMD-22671; Structure I-FS: PDBID:7K53 EMDB:EMD-22672; Structure II-FS: PDBID:7K54 EMDB:EMD-22673; Structure III-FS: PDBID:7K55 EMDB:EMD-22674.

## ACKNOWLEDGEMENTS

We thank Chen Xu and Kangkang Song for grid screening and data collection at the cryo-EM facility at UMass Medical School; and members of the Korostelev laboratory for discussions and comments on the manuscript. This study was supported by LL2008 project, MEYS CR as a part of the ERC CZ program and by Czech Science Foundation, project no. 20-16013Y (to G.D.), by NIH Grants R35 GM134931 (to Y.M.H.) and R35 GM127094 (to A.A.K.).

## AUTHOR CONTRIBUTIONS

Conceptualization: Y.M.H, A.A.K. Methodology: G.D., A.B.L., H.G, Y.M.H, A.A.K. Validation: G.D, A.A.K. Investigation: G.D., A.B.L., E.S. Resources: Y.M.H, A.A.K. Writing-Original Draft: G.D., A.A.K. Writing-Review and Editing: All; Visualization: G.D. Supervision: A.A.K. Funding Acquisition: G.D, Y.M.H, A.A.K.

## SUPPLEMENTARY TABLES AND FIGURES

**Table S1.** Refinement statistics for cryo-EM structures of non-frameshifting and frameshifting complexes.

|  | I | II | III | I-FS | II-FS | III-FS |
| --- | --- | --- | --- | --- | --- | --- |
| PDB | 7K50 | 7K51 | 7K52 | 7K53 | 7K54 | 7K55 |
| EMDB | EMD- | EMD- | EMD- | EMD- | EMD- | EMD- |
|  | 22669 | 22670 | 22671 | 22672 | 22673 | 22674 |
| **Data collection and processing** |  |  |  |  |  |  |
| Magnification | 130000x | 130000x | 130000x | 130000x | 130000x | 130000x |
| Voltage (kV) | 300 | 300 | 300 | 300 | 300 | 300 |
| Electron exposure (e<sup>-</sup>/Å<sup>2</sup>) | 47.5 | 47.5 | 47.5 | 47.5 | 47.5 | 47.5 |
| Defocus range (μm) | 0.8-2.0 | 0.8-2.0 | 0.8-2.0 | 0.8-2.0 | 0.8-2.0 | 0.8-2.0 |
| Pixel size (Å) | 0.525 | 0.525 | 0.525 | 0.525 | 0.525 | 0.525 |
| Symmetry imposed | C1 | C1 | C1 | C1 | C1 | C1 |
| Initial particle images (no.) | 62,716 | 62,716 | 62,716 | 164,504 | 164,504 | 164,504 |
| Final particle images (no.) | 4,263 | 3,179 | 4,612 | 12,108 | 9,059 | 6,029 |
| Map resolution (Å)<sup>***</sup> | 3.4 | 3.5 | 3.4 | 3.2 | 3.2 | 3.3 |
| FSC threshold |  |  |  |  |  |  |
| **Refinement** |  |  |  |  |  |  |
| Initial model used (PDB code) | 5UYM | 5UYM | 5UYM | 5UYM | 5UYM | 5UYM |
| Model resolution (Å) | 3.2 | 3.2 | 3.2 | 3.2 | 3.2 | 3.2 |
| Correlation Coefficient (cc_mask)<sup>*</sup> | 0.829 | 0.791 | 0.807 | 0.842 | 0.815 | 0.800 |
| Real space R-factor <sup>†</sup> | 0.224 | 0.250 | 0.242 | 0.224 | 0.244 | 0.256 |
| Map sharpening *B* factor (Å<sup>2</sup>) | -80 | -80 | -80 | -80 | -80 | -80 |
| Model composition* |  |  |  |  |  |  |
| Non-hydrogen atoms | 149700 | 153371 | 153371 | 149698 | 153369 | 153369 |
| Protein residues | 5944 | 6629 | 6629 | 5944 | 6629 | 6629 |
| RNA residues | 4811 | 4732 | 4732 | 4811 | 4732 | 4732 |
| Mg<sup>2+</sup>/GDPCP | 0/0 | 1/1 | 1/1 | 0/0 | 1/1 | 1/1 |
| *B* factors (Å<sup>2</sup>)* |  |  |  |  |  |  |
| Protein | 146.4 | 165.9 | 169.4 | 139.6 | 145.1 | 152.5 |
| RNA | 146.4 | 152.0 | 161.1 | 135.1 | 133.1 | 144.1 |
| R.m.s. deviations*.§ |  |  |  |  |  |  |
| Bond lengths (Å) | 0.010 | 0.012 | 0.011 | 0.009 | 0.011 | 0.012 |
| Bond angles (°) | 1.125 | 1.252 | 1.224 | 1.100 | 1.170 | 1.261 |
| Validation** |  |  |  |  |  |  |
| MolProbity score | 2.64 | 2.84 | 2.73 | 2.55 | 2.68 | 2.81 |
| Clashscore | 19.22 | 25.11 | 21.37 | 17.92 | 21.23 | 22.90 |
| Poor rotamers (%) | 1.87 | 2.08 | 1.88 | 1.52 | 1.79 | 2.19 |
| Ramachandran plot** |  |  |  |  |  |  |
| Favored (%) | 85.09 | 81.68 | 82.71 | 84.77 | 84.36 | 82.67 |
| Allowed (%) | 13.21 | 16.48 | 15.89 | 14.08 | 14.06 | 15.80 |
| Disallowed (%) | 1.70 | 1.84 | 1.40 | 1.15 | 1.58 | 1.53 |
| Validation (RNA)** |  |  |  |  |  |  |
| Good sugar pucker (%) | 99.4 | 99.5 | 99.5 | 99.4 | 99.2 | 99.5 |
| Good backbone (%) | 87.5 | 85.5 | 88.1 | 87.9 | 86.8 | 86.4 |

\*\*\* from Frealign (FSC\_part)
\*\* from Molprobity
\* from Phenix
† from RSRef
§ root mean square deviations

**Supplementary Figure 1.**
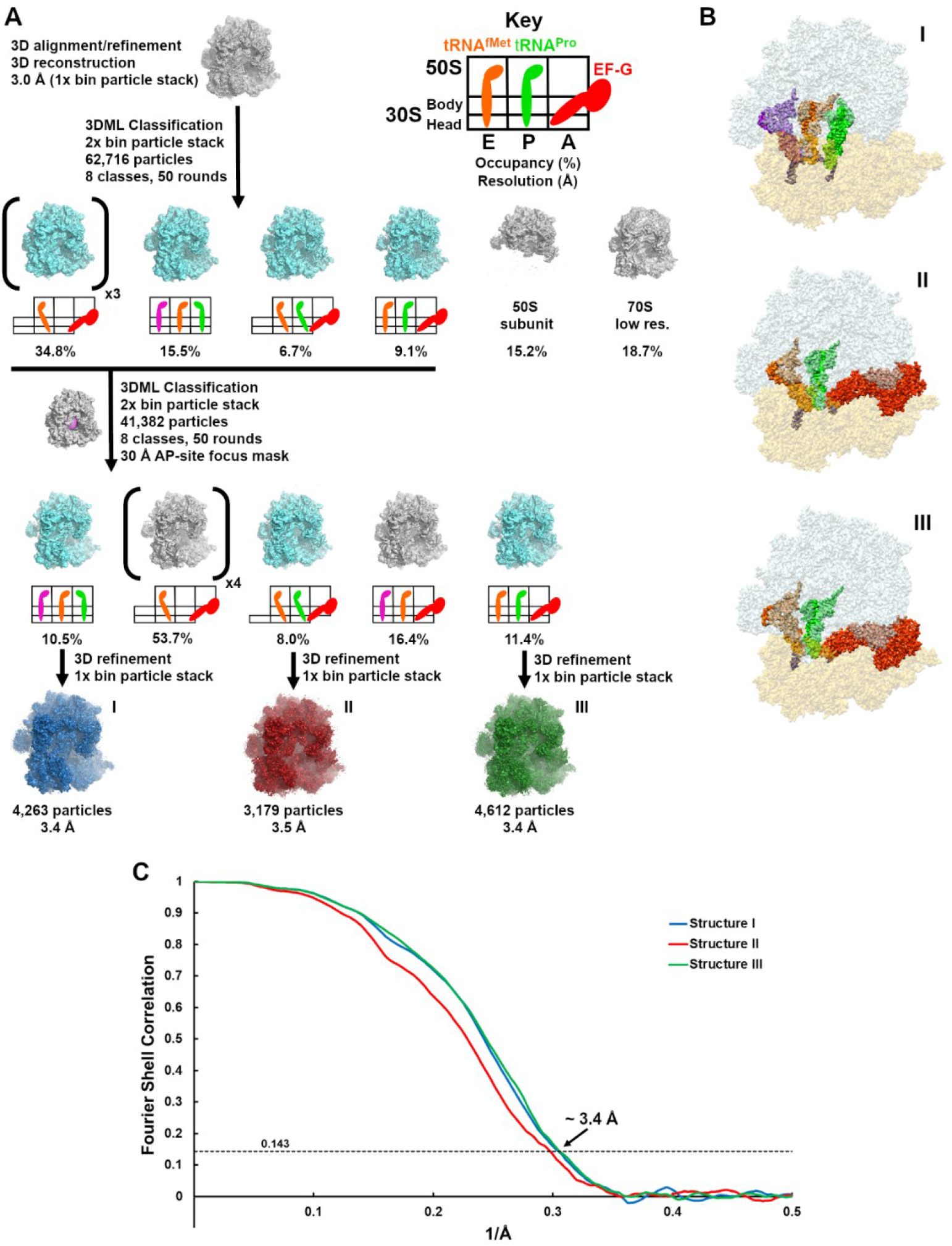
Scheme of maximum-likelihood classification resulting in cryo-EM maps of 70S ribosomes bound with and without EF-G for the non-frameshifting complex. (**A**) Classification of the dataset obtained for 70S ribosomes with the non-frameshifting CCA-A mRNA. (**B**) Segmented cryo-EM maps corresponding to Structures I, II, and III. The maps are colored as in Figure 1. (**C**) FSC between even- and odd-particle half maps for the pre-translocation and EF-G-bound translocation non-frameshifting complexes.

**Supplementary Figure 2.**
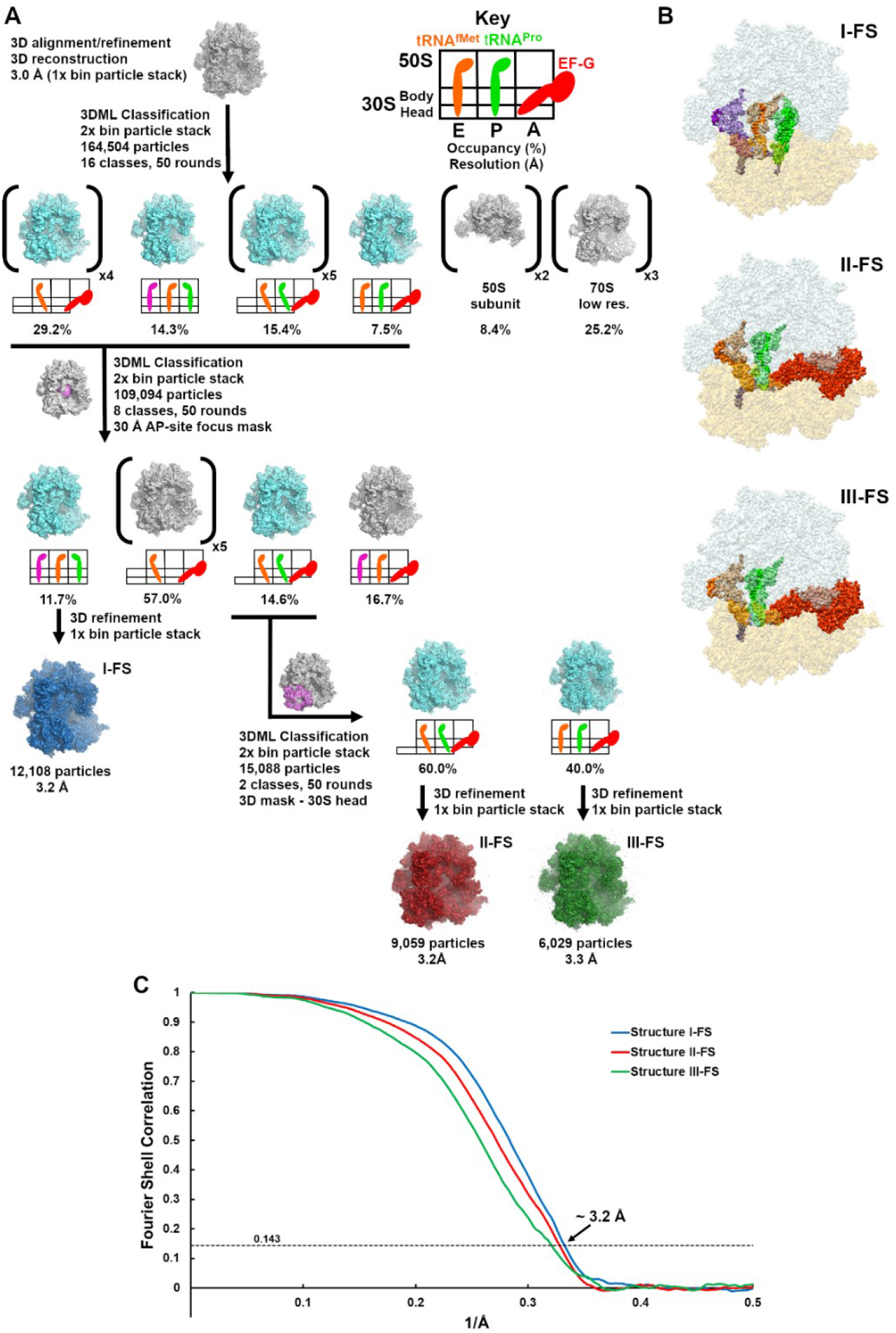
Scheme of maximum-likelihood classification resulting in cryo-EM maps of 70S ribosomes bound with and without EF-G for the frameshifting complex. (**A**) Classification of the dataset obtained for 70S ribosomes with the frameshifting CCC-A mRNA. (**B**) Segmented cryo-EM maps corresponding to Structures I-FS, II-FS, and III-FS. The maps are colored as in Figure 1. (**C**) FSC between even- and odd-particle half maps for the pre-translocation and EF-G-bound translocation frameshifting complexes.

**Supplementary Figure 3.**
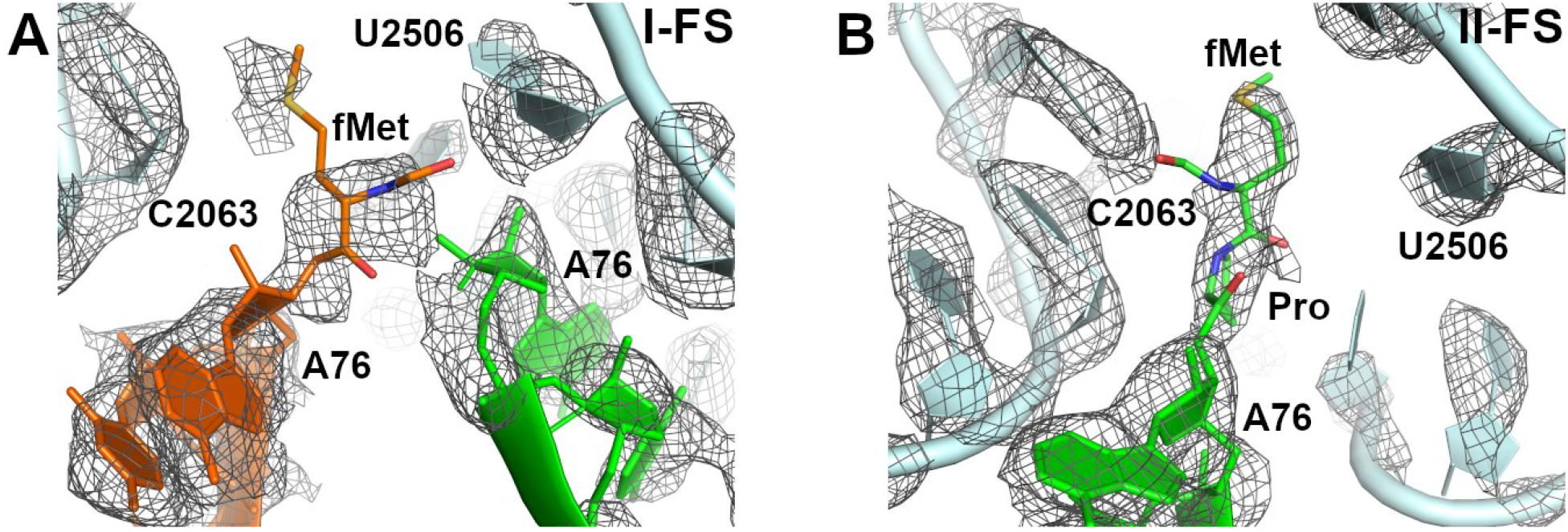
Cryo-EM density (mesh) of the peptidyl transferase center of the frame-shifting structures in the pre-translocation (**A**, I-FS) and EF-G-bound translocation (**B**, II-FS) states. In the pre-translocation state (in both Structure I and I-FS, shown in panel A), density does not allow unambiguous interpretation of the peptidyl-transfer states of the amino acids fMet and Pro, suggesting a mixture of aminoacyl- and dipeptidyl-tRNA states. fMet was modeled in both structures, because continuous density is observed between the P-tRNA nucleotide A76 and the amino-acyl moiety. The maps were sharpened by applying the B-factor of -80 Å^2^ and are shown at 2.5 σ.

**Supplementary Figure 4.**
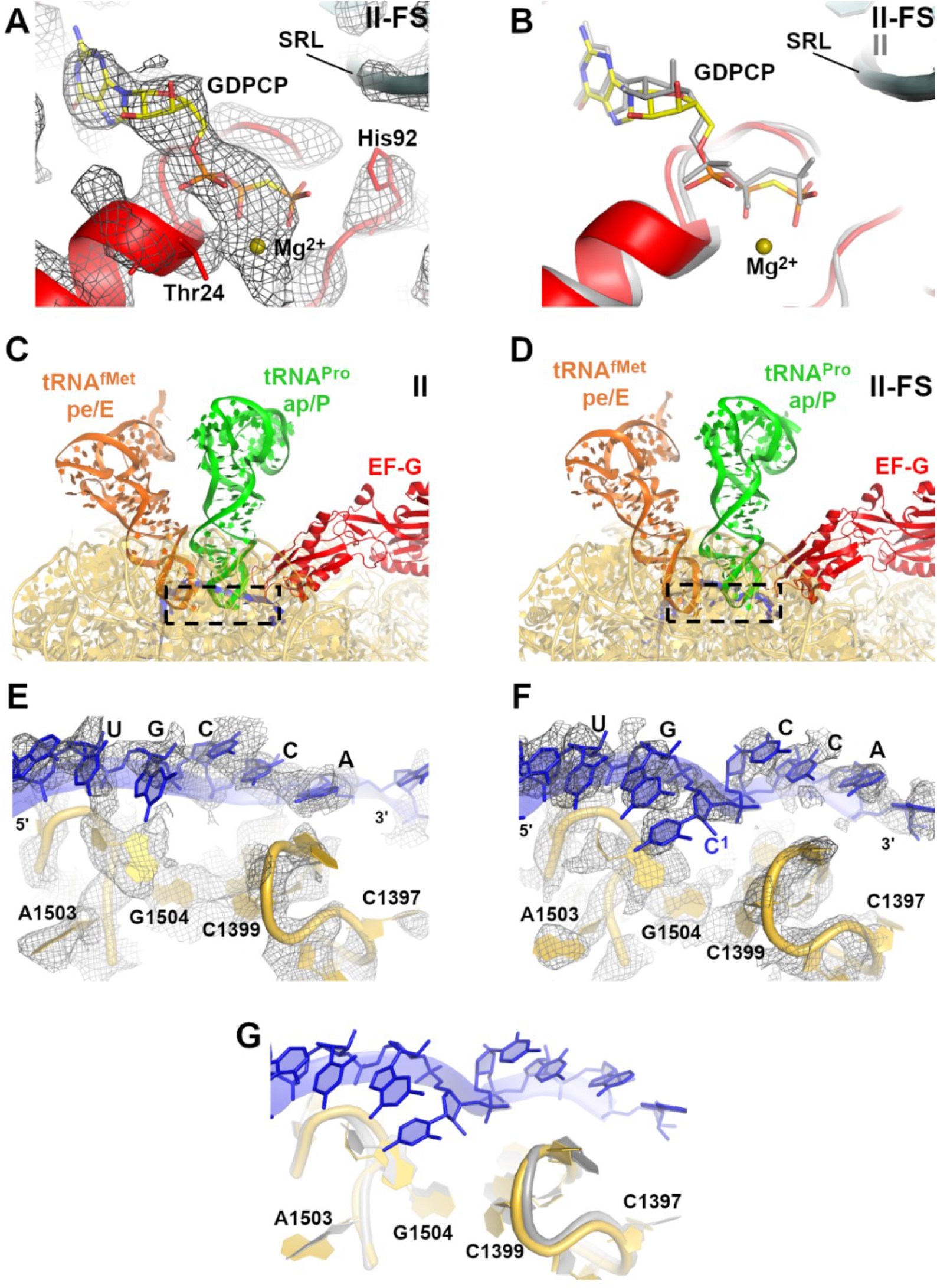
The GTPase region of EF-G and 16S rRNA surrounding the E and A sites in the EF-G-bound translocation complexes. (**A**) Cryo-EM density of the EF-G GTPase center with GDPCP in structure II-FS. The map is sharpened by applying the B-factor of -80 Å^2^ and is shown at 2.5 σ. (**B**) Structural alignment of the GTPase center (red, II-FS) and GDPCP (yellow, II-FS) with those of the non-frameshifted structure (gray, II). Structural alignment was performed by superposition of 23S rRNAs. (**C-G**) Comparison of mRNA-binding regions of the 16S rRNA between Structures II and II-FS; cryo-EM densities are shown in panels E and F, alignment of structures II-FS and II (gray) is shown in panel G. This comparison shows similar positions of the region containing nucleotides C1397 and A1503 previously proposed to participate in mRNA frame maintenance (see Results). Structural alignment was performed by superposition of 16S rRNAs.

